# Irradiation and nitrogen metabolism: differential responses in high yield indica and japonica rice commercial cultivars

**DOI:** 10.64898/2026.03.29.715102

**Authors:** Gastón Quero, Pedro Silva, Maria Martha Sainz, Sebastián Fernández, Sebastián Simondi, Jesus Castillo, Omar Borsani

**Affiliations:** Departamento de Biología Vegetal, Facultad de Agronomía, Universidad de la República, Garzón 809, Montevideo, Uruguay; Instituto de Ingeniería Eléctrica, Facultad de Ingeniería, Universidad de la República, Julio Herrera y Reissig 565, Montevideo, Uruguay; Area de Matemática, Facultad de Ciencias Exactas y Naturales, Universidad Nacional de Cuyo (FCEN-UNCuyo), Padre Contreras 1300, Mendoza, Argentina; Instituto Nacional de Investigación Agropecuaria (INIA), Treinta y Tres, Uruguay

**Keywords:** Nitrogen metabolism, amino acids, photosynthesis, carbon–nitrogen interaction, rice cultivar

## Abstract

Photosynthesis accounts for most of the final grain yield in rice, making improvements in radiation use efficiency (RUE) a key strategy for enhancing productivity. Agronomically, RUE is defined as the biomass produced per unit of total solar radiation or photosynthetically active radiation intercepted by the canopy. However, the interaction between carbon and nitrogen metabolism plays a critical role in determining plant growth and grain yield. Assimilated nitrogen is required for the synthesis of photosynthetic pigments and enzymes, while the reduction of nitrate (NOLL) and nitrite (NOLL), as well as the assimilation of ammonium (NHLL), depend on the reducing power and carbon skeletons generated by photosynthesis. In this study, two high-yielding rice (*Oryza sativa*) cultivars—an indica-type (El Paso 144) and a japonica-type (INIA Parao) were subjected to two nitrogen treatments (3 mM and 9 mM NOLL/NHLL) and two light intensities (850 and 1500 μmol mL² sL¹). A strong interaction between light intensity and nitrogen metabolism was observed, with contrasting responses between subspecies. These differences reflect a coordinated regulation of carbon assimilation and primary nitrogen metabolism. The results provide new insights into the metabolic strategies underlying nitrogen compound accumulation under variable irradiance. Such knowledge is essential for improving nitrogen fertilizer use efficiency and yield performance in elite rice genotypes cultivated under commercial field conditions.

## 1. Introduction

Rice (*Oryza sativa L.*), with 800 million tons produced in 2023, is the second most widely produced cereal, surpassed only by corn. The leading rice producers are in Southeast Asia, China, India, Bangladesh, Indonesia, Thailand and Vietnam, representing 75% of total rice production (Food and of the United Nations, 2025 with major processing by Our World in Data). Cultivated rice is classified into two major subspecies, indica and japonica, which diverged through independent domestication events in India and China, respectively (Gross and Zhao, 2014). Although rice yields have steadily increased over past decades, the rate of yield growth has slowed from 2.5% per year between 1962–1979 to 1.4% between 1980–2011, highlighting the increasing challenges of sustaining productivity and meeting future demand (Grassini et al., 2013; Ray et al., 2013).

Since approximately 90% of rice grain yield originates from photosynthetic carbon fixation (Xie et al., 2011), enhancing radiation use efficiency (RUE) is a major goal for improving productivity. Agronomically, RUE is defined as the biomass produced per unit of total solar or photosynthetically active radiation intercepted by the canopy (Stockle and Kemanian, 2009). The harvest index (HI), defined as the ratio of harvested grain to total plant biomass, is also a key determinant of yield potential. However, as the HI in modern cultivars approaches its theoretical maximum, further gains cannot be achieved solely by modifying plant architecture (Mitchell and Sheehy, 2006). Thus, advancing knowledge of the physiological processes underlying photosynthesis and biomass partitioning remains central to sustaining future yield improvements in rice cultivation.

Nitrogen (N) metabolism plays a central role in determining potential yield and is tightly coupled to carbon metabolism. Nitrogen is a fundamental element for all living organisms; it is required for the synthesis of amino acids, proteins, nucleic acids, and chlorophyll (Basuchaudhuri, 2016). In plants, N is mainly absorbed from the soil as nitrate (NOLL) or ammonium (NHLL). Because N is often the most limiting nutrient for crop growth, cereals tend to exhibit relatively low nitrogen use efficiency (NUE), with substantial losses of applied N through volatilization, leaching, or denitrification. In rice, fertilizer recovery efficiency rarely exceeds 30–45%, depending on management intensity (Chauhan Khawar et al., 2017; Dobermann, 2007; Ladha et al., 2020). The resulting imbalance between high fertilizer input and low recovery efficiency has critical environmental consequences (You et al., 2023).

From an agronomic perspective, NUE is defined as the yield obtained per unit of available or applied N (Pathak et al., 2008). Although breeding and management advances have significantly improved NUE in crops such as maize and rice (O’Neill et al., 2004), further progress requires a deeper molecular understanding of N uptake, assimilation, and remobilization (Kant et al., 2011). Nitrogen and carbon assimilation are strongly interconnected: the reduction of nitrate and assimilation of ammonium depend on the reducing power and carbon skeletons generated through photosynthesis, while nitrogen-derived metabolites are essential for synthesizing photosynthetic pigments and enzymes (Foyer et al., 2001).

The GS/GOGAT cycle exemplifies the interdependence of carbon and nitrogen metabolism, as ammonium assimilation requires ATP, reducing equivalents, and carbon skeletons. 2-oxoglutarate (2-OG), derived from the tricarboxylic acid (TCA) cycle, is the primary carbon skeleton consumed during glutamate synthesis. Because 2-OG is continuously depleted, replenishment of TCA intermediates is essential. Phosphoenolpyruvate carboxylase (PEPC) provides oxaloacetate (OAA), which serves both as an anaplerotic substrate for the TCA cycle and as a precursor for amino acids such as aspartate (Asp) (Masumoto et al., 2010; Monza and Marquez, 2004). This integration highlights how nitrogen assimilation depends on carbon fluxes generated through photosynthesis and central metabolism.

Although the coupling between C and N metabolism has been extensively studied, the influence of light irradiance on nitrogen assimilation and its downstream effects on biochemical and physiological traits in modern cultivars remains underexplored. Most previous studies were conducted in tropical or subtropical regions, where solar radiation is consistently high (Makino, 2011; Peng et al., 2004). In contrast, temperate rice systems, such as those in southern South America, experience more variable irradiance and cooler temperature factors that may significantly alter the coordination between photosynthesis and nitrogen assimilation. Recent research emphasizing improvements in canopy photosynthesis underscores the need to examine these processes under temperate conditions to develop genotype- and environment-specific strategies (Vishwakarma et al., 2023; Xiong, 2024).

This study hypothesizes that light irradiance differentially regulates nitrogen metabolism in high-yielding *indica* and *japonica* rice cultivars adapted to temperate conditions. These differences can be elucidated by examining biochemical and physiological parameters associated with nitrogen assimilation and utilization.

## 2. Materials and Methods

### 2.1 Plant material and growth conditions

Two high-yielding rice cultivars (*Oryza sativa L.*) were used as plant material: the indica-type cultivar El Paso 144 and japonica-type cultivar INIA Parao, both with an estimated yield potential of around 14 Mg ha^-1^ (Carracelas et al., 2023).

The seeds germinated for five days in Petri dishes partially covered by distilled H_2_O in the dark at 28 °C. The germinated plants were transferred to growth supports, which consisted of PVC tubes with a volume of 850 cm^3^ containing vermiculite sand substrate 1:1 (v/v). The plants grew on these supports for 40 days in growth chambers with the following conditions: photoperiod was 12/12h (light/dark), temperature 26-30 °C and relative humidity of 60-80%. Two light regimes were applied: LTI, 850 μmol mL² sL¹ (representing half a full-sun day), and LTII, 1500 μmol mL² sL¹ (simulating full sunlight). LED luminaires providing a continuous spectrum between 400–700 nm were used (Quero et al., 2019; Supplementary Fig. 1S). The photoperiod reproduced the natural light curve of a sunny day in a rice-growing region of Uruguay (33°13′51″S, 54°22′56″W).

Plants were irrigated weekly with 100 mL of a modified Yoshida nutrient solution (Yoshida et al., 1971; Supplementary Table 1S) adjusted to pH 5.2. The nitrogen source was NHLNOL supplied at two concentrations: N3, 1.5 mM (representing the standard commercial fertilization level), and N9, 4.5 mM (high nitrogen). Additional watering with distilled water ensured adequate moisture throughout the growth period.

### 2.2 Nitrogen and carbon content

Shoots were harvested after 40 days of growth, oven-dried at 50 °C to constant weight, and ground to a fine powder using an agate mortar. Total carbon and nitrogen concentrations were determined at the Center for Applications of Nuclear Technology in Sustainable Agriculture (Faculty of Agronomy, Universidad de la República, Uruguay) using an elemental analyzer.

### 2.3 Photosynthetic parameters

#### Energy partitioning: quenching and relaxation analyses

The quantification of energy partition in PSII was determined by the quantum yield of three de-excitation processes using the chlorophyll’s fluorescence parameters (Lazár, 2015; Quero et al., 2021). A quenching analysis (light-adapted plants) was performed for this purpose, followed by a relaxation analysis (dark-adapted plants after light period) of PSII as proposed by Kasajima et al. (2009) and revised by Lazár (2015). This analysis is based on the idea that the sum of all the de-excitation processes of the energy absorbed by PSII is equal to 1 (Demmig-Adams et al., 1996; Hendrickson et al., 2004; Kramer et al., 2004; Logan et al., 2014):

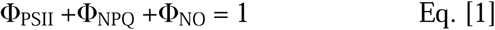

The quenching analysis of Φ_PSII_ was calculated as Eq. [1] (Table 1), (Baker, 2008; Genty et al., 1989). Φ_NPQ_ and Φ_NO_ were calculated as Eq. [2] and [6], respectively (Table 1)(Hendrickson et al., 2004; Klughammer and Schreiber, 2008). In the relaxation analysis of PSII, the quantum yields of the fast and slow components of the non-photochemical quenching Φ_NPQf_ and Φ_NPQs_ were calculated as Eq. [4] and Eq. [5], respectively (Table 1) (Kasajima et al., 2009) Chlorophyll fluorescence analysis was measured with an amplitude-modulated fluorescence monitor (FMS1, Hansatech®). The plants were measured in darkness after one hour of adaptation. First, a fluorescence quenching analysis was carried out, followed by a relaxation analysis as presented in Supplementary Fig. 2S (Quero et al., 2019).

**Table 1.**
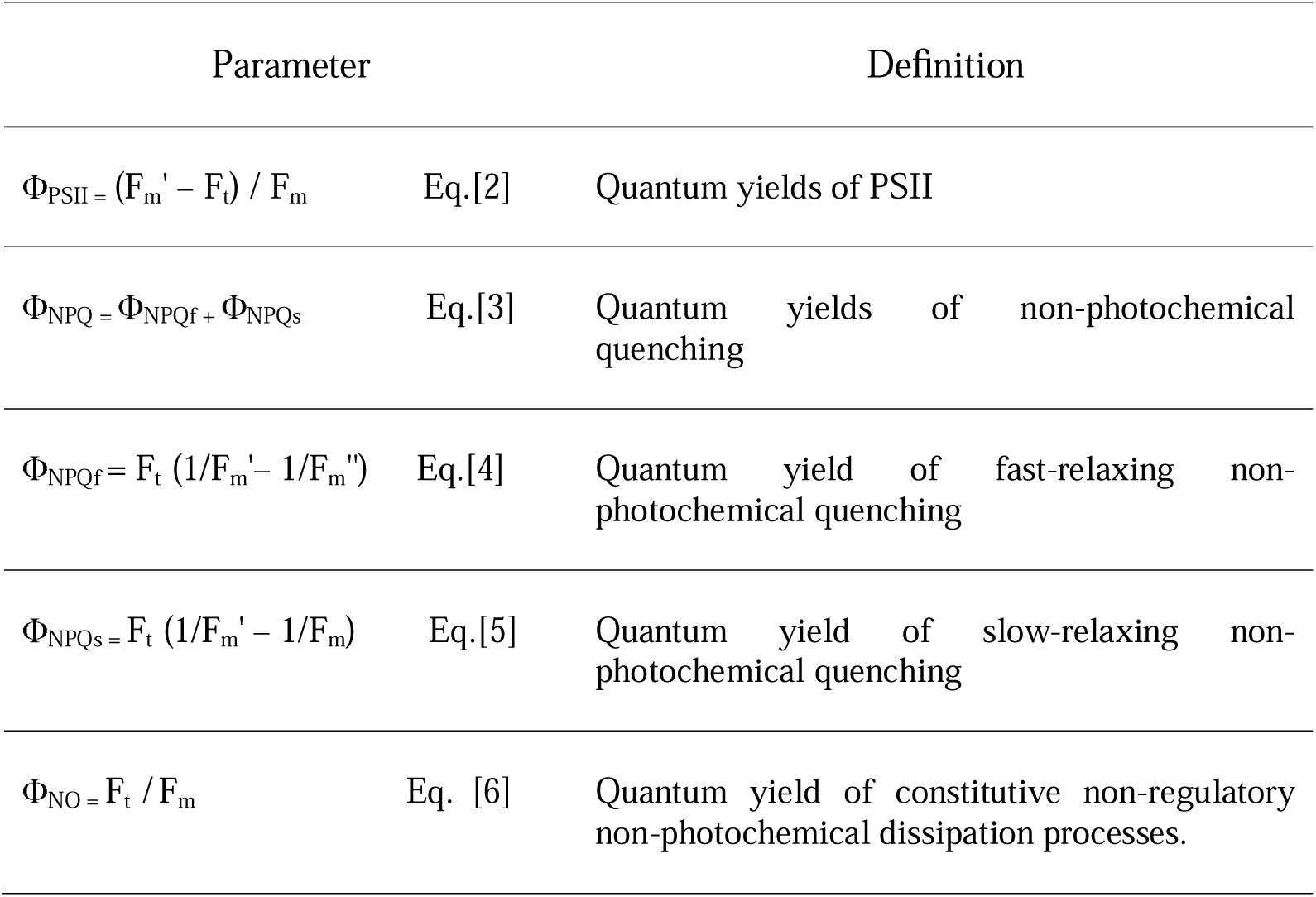
Quantum yields parameter and determining equations.

#### Carbon assimilation

CO_2_ assimilation rate and chlorophyll fluorescence analyses were performed on the newest expanded leaf. CO_2_ assimilation was measured with a portable photosynthesis system (LI-6400, LI-COR®). Photosynthesis measurements were carried out with the plants after three hours of light exposure. The CO_2_ concentration was 395 ± 5 ppm, the humidity was 60%, and the temperature was 25 °C. The intracellular concentration of CO_2_, the transpiration rate, stomatal conductance, and net photosynthesis were measured with an incident photon flux equal to that of the conditions in which the plants grew, 1500 μmol m^-2^ sec^-1^ or 850 μmol m^-2^ sec^-1^ for high and low radiation, respectively.

### 2.4 Protein Extraction and Quantification

Proteins were extracted according to Li et al. (2011) 0.1 g of previously ground leaves were macerated with liquid nitrogen, with 0.5 ml of 62.5 mM TRIS-HCl buffer pH 7.4; 2 mM EDTA; 2 % (v/v) 2-mercaptoethanol; 1.0 mM phenylmethylsulfonyl fluoride (PMSF); 0.1 % sodium dodecyl sulfate (SDS); 10 % glycerol. The homogenate was centrifuged at 15000 x g for 15 min at 4 °C. The supernatant was used as a crude extract for the quantification of soluble proteins and for immunodetection assays.

The extractions for the enzymatic activity assays were performed by changing the extraction buffer for 100 mM TRIS-HCl pH 7.5; MgCl2 10 mM; dithiothreitol 1.0 mM; PMSF 1.0 mM; Triton x-100 0.1 % (v/v). The rest of the extraction was the same as that performed for the immunodetection assays. Proteins were determined by the Bradford method (1976). 980 μL of reagent consisting of 0.1 mM Coomassie Blue G250 (Sigma), 0.8 M ethanol and 1.5 M phosphoric acid were added to 20 μL of sample. Absorbance was measured at 595 nm after 2 min; known concentrations of Lysozyme were used for the standard curve.

### 2.5 Nitrogen compound: amino acids, ammonium and nitrate quantification

Total amino acids, ammonium, and nitrate were extracted according to Izaguirre-Mayoral et al. (1992). HPLC used the supernatant to quantify ammonium, nitrate, and total amino acids and analyze amino acids and nitrogenous compounds. Total amino acids were determined according to Moore and Stein (1948). Nitrate was quantified according to Cataldo et al. (1975) with modifications. To 0.1 mL of the sample, 0.4 mL of 0.36 M salicylic acid in 18.4 M sulfuric acid was added, and the sample was shaken. After 20 minutes, 5.0 mL of 3.8 M NaOH was added. The absorbance was measured at 410 nm, and KNO_3_ was used as a standard. Ammonium was quantified according to Solorzano (1969).

The amino acids were identified and quantified after derivatization with ophthaldialdehyde (OPA, Sigma-Aldrich) according to Hunkapiller et al. (1984) with modifications. To 20.0 μL of extract, 40.0 μL of 0.4 M borate-NaOH buffer pH 10.0; 0.7 M 2-mercaptoethanol and 86.0 μM OPA in methanol-water (1/1) were added. The samples were shaken for 4 min at 2500 rpm, centrifuged at 12000 x g for 2 min, and injected into an HPLC equipped with a 20.0 μL unloop. The elution and separation of the amino acids were carried out using a C18 column as the stationary phase, and two solvents were used as the mobile phase. The first solvent was 20.0 mM acetate buffer pH 5.5 (solvent A), and the second was methanol (solvent B). The gradient elution condition, at a 1.0 mL/min flow and 25 °C. Thus, a calibration curve was determined for each amino acid-OPA.

### 2.6 Immunodetection of GS and Fd-GOGAT

For the immunodetection of enzymes, total protein electrophoresis under denaturing conditions was performed on SDS-polyacrylamide gels. For the preparation of the gel and buffer, the protocol proposed by (Sambrook and Rusell, 2000) was followed. The gels were equilibrated for 10 min in transfer buffer: 25.0 mM Tris and 192.0 mM glycine pH 8.3, 20% methanol for protein transfer. Wet electro transfer was performed at 31 mA at 4 °C for 3 h. The transfer of proteins to the membrane was checked by staining with Ponceau Red 0.5% (w/v) and acetic acid 1% (v/v) until the bands appeared. For the immunodetection of the Fd-GOGAT and GS proteins, polyclonal rabbit anti-Fd-GOGAT (Agrisera, catalog number #AS07 242) and barley anti-GS antibodies were used, provided by Prof. M. Betti (Faculty of Chemistry, Universidad de Sevilla, Spain) in a 1/1000 dilution. A polyclonal goat anti-rabbit IgG antibody (Agrisera, catalog number #AS09 602), conjugated with HPR (1/10000) was used as a secondary antibody. The membranes were developed in a C-Digit® viewer.

### 2.7 Experimental design and Statistical treatment

The experiment followed a completely randomized design with three factors: genotype (El Paso 144 and INIA Parao), light irradiance (LTI and LTII), and nitrogen level (N3 and N9), resulting in eight treatment combinations. Each experimental unit consisted of one plant per pot, with three biological replicates per treatment. Data were analyzed by three-way ANOVA, and significant differences among means were determined using orthogonal contrast analysis at *P* ≤ 0.05. Statistical analyses were performed in R (R Core team, 2024) using the packages *stats*, *lme4* (Bates et al., 2015), and *emmeans* (Lenth, 2025). Pearson correlation coefficients were calculated using the base *cor* function.

## 3. Results

### 3.1 Primary nitrogen metabolism

Nitrogen concentration in leaf tissue was primarily determined by the nitrogen supply but was also significantly influenced by light intensity. Plants grown under high nitrogen (N9) exhibited higher tissue N concentrations than those under low nitrogen (N3), and this difference was amplified under high irradiance (LTII). Conversely, total carbon percentage remained unaffected by either light or nitrogen treatments (Table 2).

**Table 2.** Carbon and nitrogen content of rice leaves tissue.

| Genotype | Light | Nitrogen | N% | C% |
| --- | --- | --- | --- | --- |
| El Paso 144 | LTI | N3 | 2.1 a | 38 a |
|  |  | N9 | 3.4 c | 38 a |
|  | LTII | N3 | 2.8 b | 36 a |
|  |  | N9 | 4.2 d | 37 a |
| INIA Parao | LTI | N3 | 2.4 a | 38 a |
|  |  | N9 | 3.5 c | 38 a |
|  | LTII | N3 | 2.8 b | 37 a |
|  |  | N9 | 4.5 d | 38 a |

Same letters indicate no significative difference at $P < 0.05$

Plants were fed by a nutrient solution containing the same concentration of nitrate and ammonium (1.5 mM and 4.5 mM for N3 and N9, respectively), to analyze the impact of light irradiance on assimilation of these inorganic N forms the content of both ions was analyzed. In this study, the ammonium was undetectable, indicating the preference to assimilate this N form; however, the nitrate concentration in the green tissues varied according to genotype and light regime. Both genotypes accumulated more nitrate with increasing nitrogen supply and irradiance, though the magnitude differed between subspecies (Fig. 1). In El Paso 144 (indica), nitrate content increased approximately sixfold from N3 to N9 under high light. In INIA Parao (japonica), this increase was much greater, rising from twofold under low light (LTI) to over thirteenfold under high light (LTII). Ammonium levels remained undetectable, indicating rapid assimilation of this nitrogen form.

**Fig. 1.**
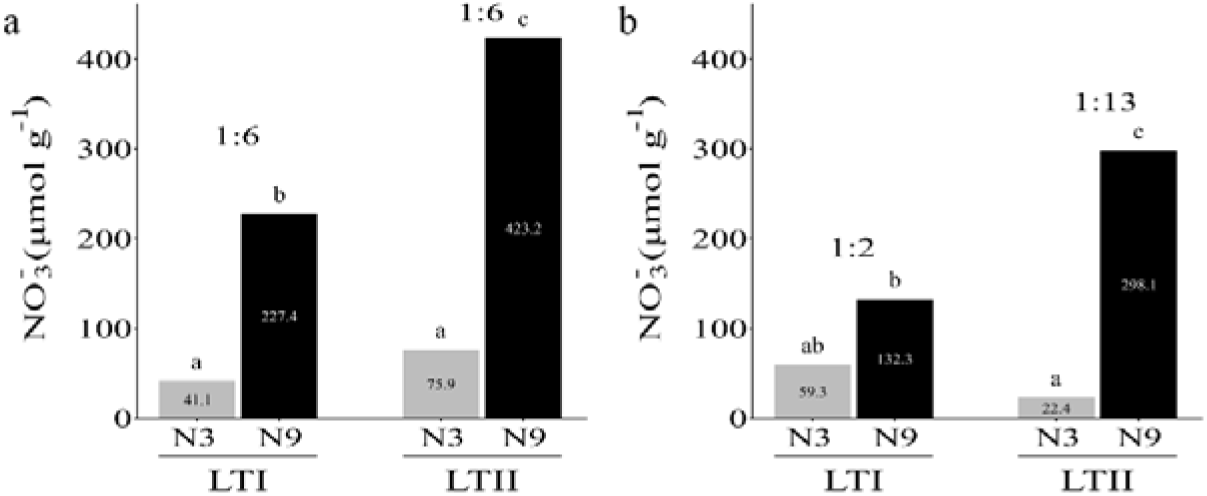
Nitrate (NO_3_^-^) content in rice leaves. N3 and N9 indicate plants grown under 3 and 9 mM of NO_3_NH_4,_ respectively. LTI and LTII refer to light irradiation levels of 850 µmol m^-2^ s^-1^ and 1500 µmol m^-2^ s^-1^ respectively. a) El Paso 144 (indica cultivar), b) INIA Parao (japonica cultivar). Numbers above the bars indicate the ratio between nitrate content in the N3 treatment and that in N9 at the same irradiation level. Bars with the same letters are not significative different at P < 0.05.

Amino acids were the first molecules derived from the primary assimilation of a nitrogen source (nitrate and ammonium) and have as the main destination the protein to which most of the assimilated N is destined. The total content of amino acids was measured in the leaves of the plants grown in the different treatments to analyze their availability for protein synthesis. The concentration of amino acids in the leaves depends mainly on the light radiation in which the plants grow. Plants grown in low radiation (LTI) presented a higher accumulation of amino acids (Fig. 2). In the case of plants grown in high radiation (LTII), El Paso 144 cultivar accumulated similar amino acids content of LTI treatment only when plants were fed with high dose of N (Fig. 2a), however INIA Parao cultivar accumulated less amino acid content in LTII than LTI treatment in both nitrogen regimes (Fig. 2b).

**Fig. 2.**
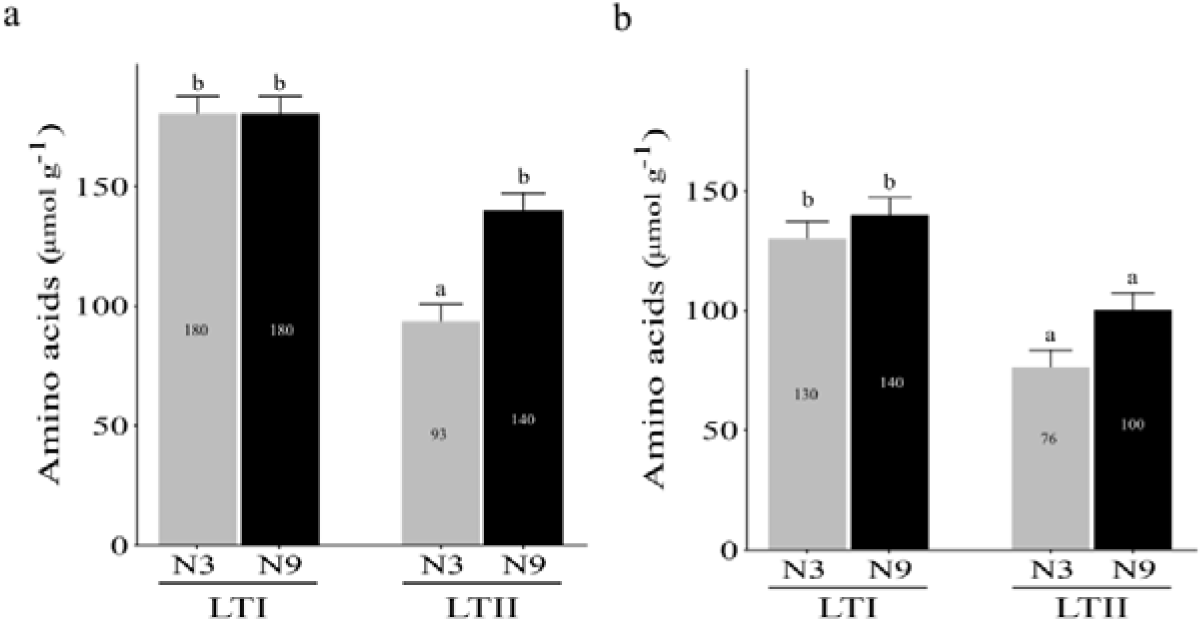
Free amino acids form rice leaves. N3 and N9 indicate plants grown under 3 and 9 mM of NO_3_NH_4_, respectively. LTI and LTII refer to light irradiance levels of 850 µmol m^-2^ s^-1^ and 1500 µmol m^-2^ s^-1^ respectively. a) El Paso 144 (indica cultivar), b) INIA Parao (japonica cultivar). Bars with the same letters are not significantly different at P < 0.05.

An analysis of the composition of the extracted and quantified amino acids was carried out to determine whether the treatments modified the relative quantities of specific amino acids. Variations in the relative amounts of any amino acid may suggest that its synthesis pathway is favored by certain growth conditions. This analysis focused on quantifying the main amino acids synthesized during assimilation and transport of amino acids. Since this complicates the analysis of the behavior of certain amino acids, it was then decided to measure the amounts of asparagine (Asn) and glutamine (Gln) with respect to the amounts of the precursor amino acids aspartate (Asp) and glutamate (Glu), respectively (Fig. 3).

**Fig. 3.**
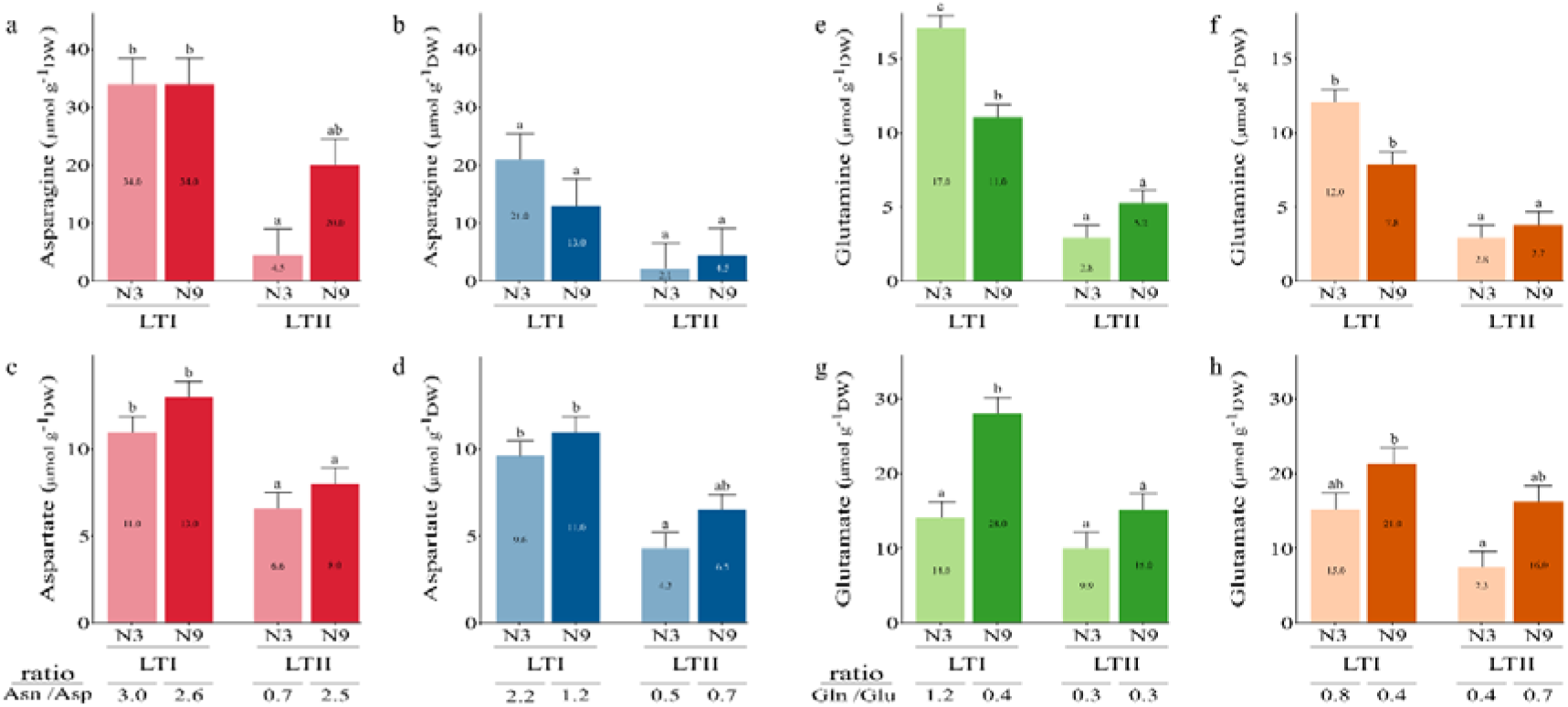
Relative contents of asparagine, aspartate and glutamine, glutamate in rice leaves. N3 and N9 indicate plants grown under 3 and 9 mM of NO_3_NH_4_, LTI and LTII refer to irradiation 850 µmol m^-2^ s^-1^ and 1500 µmol m^-2^ s^-1^ respectively. a) c) e) g) El Paso 144 (indica cultivar) b) d) f) h) INIA Parao (japonica cultivar). Asn: asparagine, Asp: aspartate, Gln: glutamine, Glu: glutamate. Same letters indicate non significative difference at P < 0.05.

The Asn/Asp ratio showed differences in the responses of the cultivars to the treatments. For indica cultivar El Paso 144, the radiation and the N dose in which the plants grew determined the amount of Asn with respect to Asp (Fig. 3 a and c). The low radiation (LTI) in both nitrogen dose and high radiation (LTII) treatments with high dose of N (N9) caused a significant increase in the Asn/Asp ratio compared to the high radiation treatment and control dose of N (N3) (Fig. 3 a and c). In the case of japonica cultivar INIA Parao, only the low radiation treatment combined with control nitrogen dose treatment generated an increase in the Asn/Asp ratio (Fig. 3 b and d). In the case of the Gln/Glu ratio, the same response was observed to radiation and N dose in both cultivars. The cultivars presented a significantly higher Gln/Glu ratio in the low radiation treatment (LTI) and control dose of N (N3) (Fig. 3 e and g). In the remaining growth conditions, there was no difference in the Gln/Glu ratio (Fig. 3).

The GS-GOGAT cycle is the main pathway for N assimilation; factors such as the dose of N and radiation under which plants grow could regulate this pathway. Furthermore, previous results regarding the Gln/Glu ratio suggest that some enzymes involved in the synthesis of these amino acids were affected by the nitrogen and light conditions. The specific activity of the GS enzyme was measured; however, no significant difference was observed between the treatment and cultivar (Supplementary Table 2S). Nevertheless, the quantity of enzyme could be affected by growth conditions. In this regard, Western blot analyses were performed to determine if there were differences in the amounts of GS and Fd-GOGAT in response to the treatments. The amount of Fd-GOGAT was affected by the N dose in which the plants were grown. Plants grown in high doses of N (N9) presented a greater amount of Fd-GOGAT regardless of the cultivar (Fig. 4). The Western blots of GS allowed us to identify the two isoforms of this enzyme, the plastid (GSp) and the cytosolic (GSc). Treatments with high doses of N (N9) caused an increase in the amount of GSp compared to GSc. Furthermore, this increase was always more significant in high radiation and high N treatment (LTII N9) compared to the low radiation treatment (LTI N9) (Fig. 4).

**Fig. 4.**
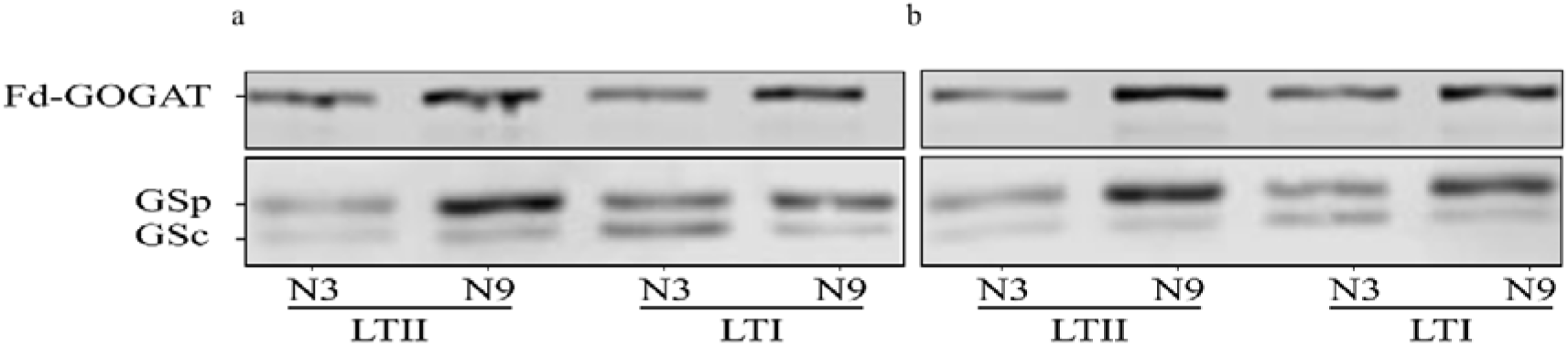
Western-blot analysis of enzymes GS and Fd-GOGAT in leaves of rice plants subjected to nitrogen and irradiation combination. N3 and N9 indicate plants grown under 3 and 9 mM of NO_3_NH_4_, LTI and LTII refer to irradiation 850 umol m^-2^ s^-1^ and 1500 umol m-^2^ s^-1^ respectively. a) El Paso 144 (indica cultivar). b) INIA Parao (japonica cultivar). GSp: Plastidic Glutamine synthetase, GSc: Cytosolic Glutamine synthetase.

### 3.2 Carbon metabolism: biochemical and photochemical phases of photosynthesis

Photosynthetic parameters were analyzed to understand how nitrogen availability interacts with carbon sources. Since the photosynthetic components involved in light energy capture, electron transport, ATP and NADPH synthesis machinery, and the enzymes of the Calvin-Benson Cycle are proteins, the combination of nitrogen and light, could affect the functionality and content of all these components. The parameters of the photochemical phase of photosynthesis appear unaffected for any combination of light or nitrogen (Table 3). On the other hand, a higher carbon fixation rate would mean more reducing power and carbon skeletons to assimilate N. In this sense, the analysis of the CO_2_ fixation rate showed that radiation was fundamental for an increase in this process. The two cultivars showed greater photosynthesis in the high radiation (LTII) treatments (Table 3), and this could be associated with a high stomatal aperture induced by this radiation level. Furthermore, the N dose in which the plants grew also affected the fixation rate. This is observed when comparing plants grown in the same radiation with different doses of N; thus, plants grown in a high dose of N tend to have a higher photosynthetic rate than the plants under a low N dose (Table 3). However, only the cultivar INIA Parao showed a significant difference in response to high light and high nitrogen (Table 3).

**Table 3.** Photosynthetic parameters in rice.

| Genotype | Light | Nitrogen | $A_n$<br>($\mu\text{mol CO}_2 \text{ m}^{-2} \text{ s}^{-1}$) | $g_w$<br>($\text{mmol H}_2\text{O m}^{-2} \text{ s}^{-1}$) | $\Phi_{\text{PSII}}$ | $\Phi_{\text{NPQf}}$ | $\Phi_{\text{NPQs}}$ | $\Phi_{\text{NO}}$ |
| --- | --- | --- | --- | --- | --- | --- | --- | --- |
| El Paso 144 | LTI | N3 | 9.9 a | 370 a | 0.34 a | 0.38 a | 0.08 a | 0.20 a |
|  |  | N9 | 10.0 a | 320 a | 0.23 a | 0.41 a | 0.09 a | 0.27 a |
|  | LTII | N3 | 11.6 a | 660 a | 0.26 a | 0.41 a | 0.07 a | 0.25 a |
|  |  | N9 | 13.7 a | 670 a | 0.32 a | 0.29 a | 0.07 a | 0.31a |
| INIA Parao | LTI | N3 | 7.4 a | 260 a | 0.23 a | 0.33 a | 0.19 a | 0.24 a |
|  |  | N9 | 11.2 ab | 630 a | 0.26 a | 0.39 a | 0.12 a | 0.22 a |
|  | LTII | N3 | 13.0 ab | 510 a | 0.31 a | 0.41 a | 0.07 a | 0.21 a |
|  |  | N9 | 16.5 b | 520 a | 0.32 a | 0.36 a | 0.08 a | 0.23 a |

Same letters indicate no significative difference at $P < 0.05$

To analyze whether differences existed in the quality of proteins accumulated by plants grown in different environments, soluble proteins were extracted from the leaves of the plants and run on denaturing polyacrylamide gels. The protein profiles depicted in Fig. 5 do not seem to have differences between them, except in the region close to 55 kDa. Based on the amount of protein observed in that band and its weight, it can be inferred that it corresponds to the largest subunit of RuBisCO (Gupta and Kim, 2015). As Fig. 5 shows, high nitrogen treatment induced an accumulation of RubisCO independently of light regime; however, this effect is more evident in the Parao cultivar. Interestingly, the higher CO_2_ assimilation observed in highlight conditions (LTII) with high nitrogen supply (N9) correlated with this possible RubisCO content.

**Fig. 5.**
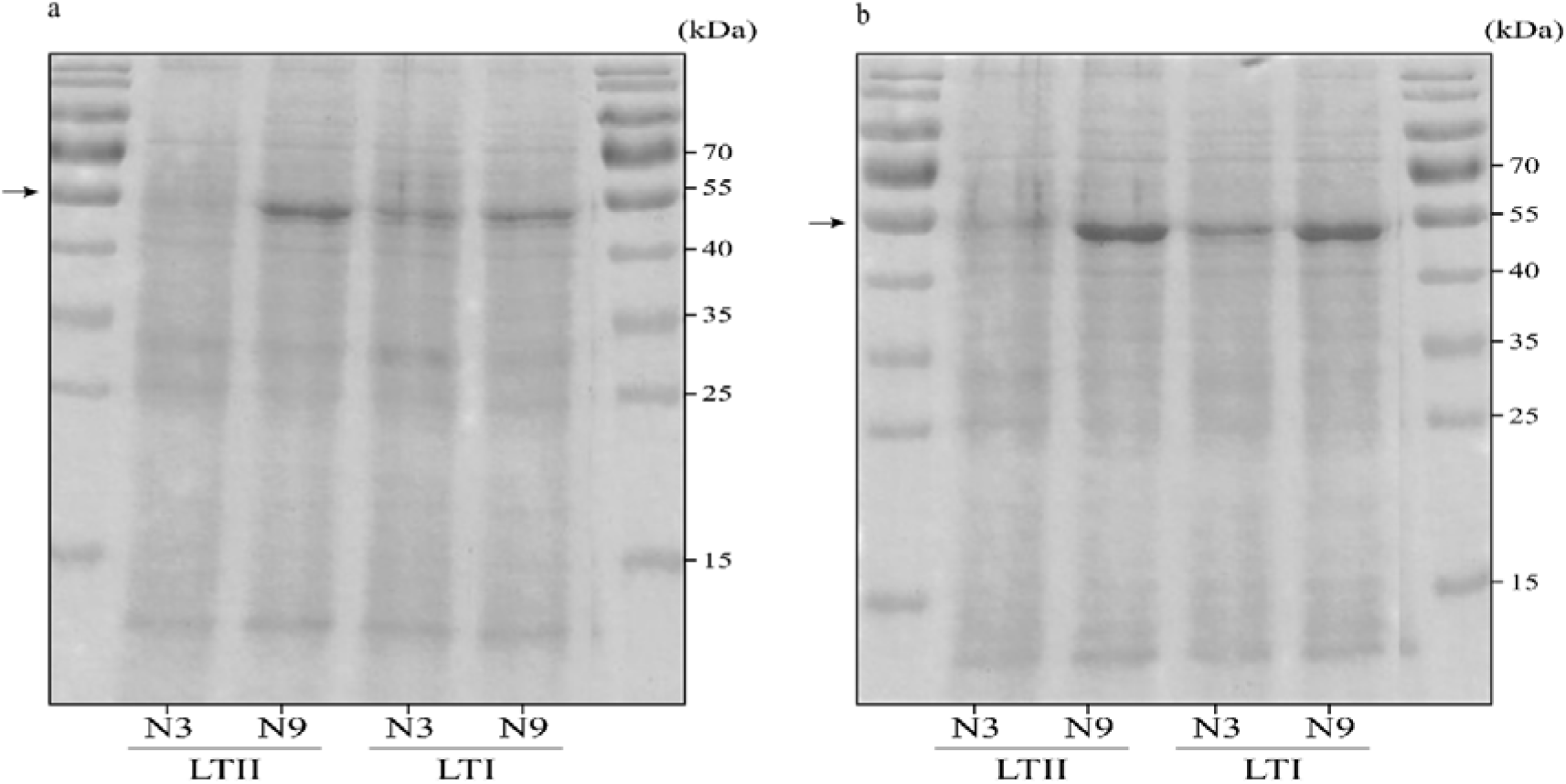
Soluble protein profile from leaves extract. Polyacrylamide denaturing gel, each line corresponds to 10 ug of protein extract of each cultivar in each treatment and stained with Coomassie blue. N3 and N9 indicate plants grown under 3 and 9 mM of NO_3_NH_4_, LTI and LTII refer to irradiation 850 µmol m^-2^ s^-1^ and 1500 µmol m^-2^ s^-1^ respectively. a) El Paso 144 (indica cultivar). b) INIA Parao (japonica cultivar). Arrows indicate the 55 kDa marker.

### 3.3 Photorespiratory-related amino acids

In addition to the amino acids related to N transport, we analyzed other amino acids that could be synthesized and accumulated differently by plants grown in different environments. The synthesis of amino acids serine and glycine is stimulated under high photorespiratory conditions (Busch et al., 2018). A good example of the connection between carbon and nitrogen metabolisms is the glycolate pathway during the photorespiration process, in which serine and glycine play an important role (Rosa-Téllez et al., 2024). Analysis of these two amino acids showed that the content is not affected by the nitrogen regimes, but the irradiation level influences the accumulation (Fig. 6). Interestingly, the lowest content in these amino acids was consistently found under high light conditions, same light conditions wherein the higher level of CO_2_ assimilation rate was observed. This was more evident in the japonica cultivar (Fig. 6).

**Fig. 6.**
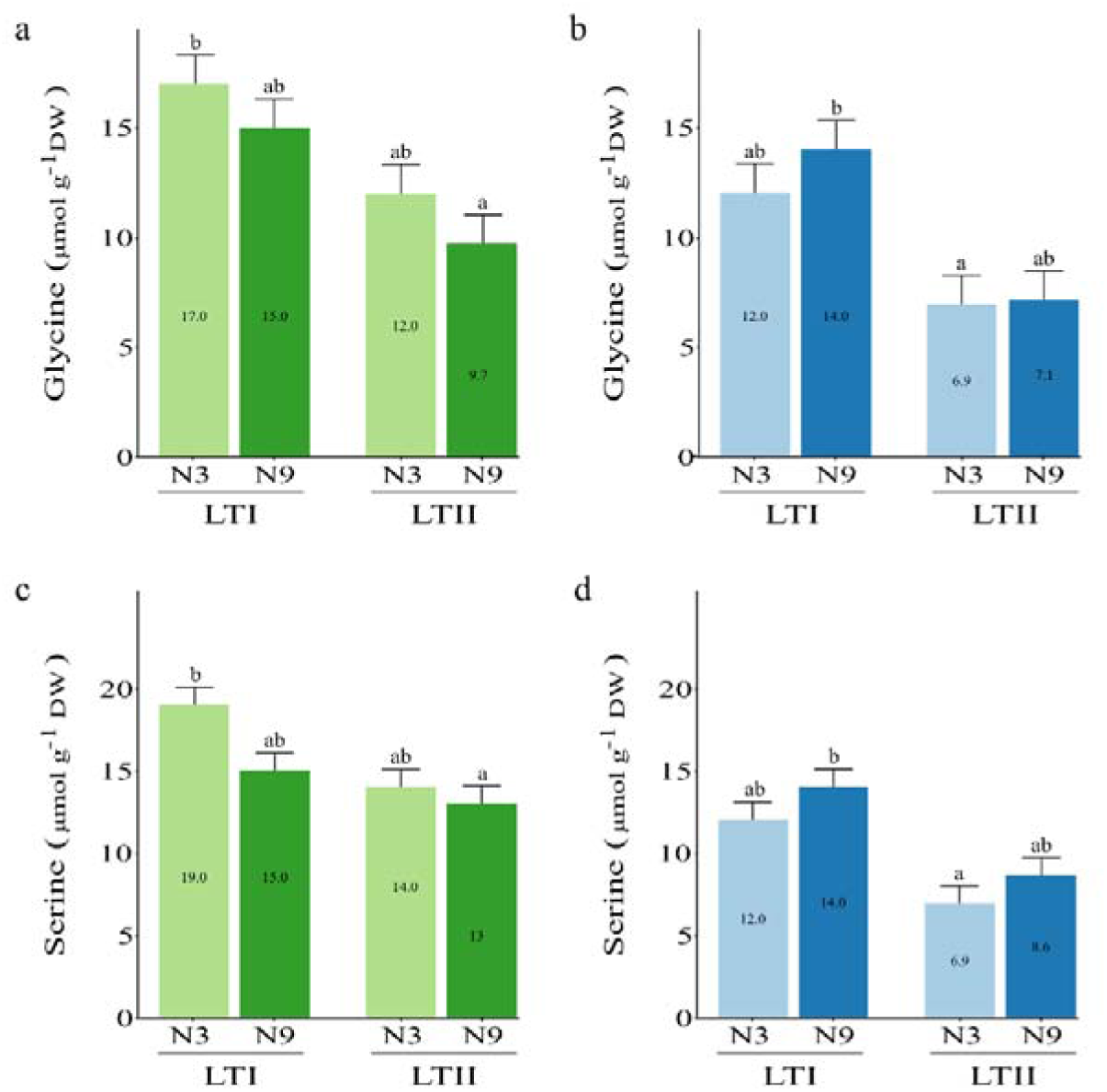
Photorespiratory amino acids glycine and serine. N3 and N9 indicate plants grown under 3 and 9 mM of NO_3_NH_4_, LTI and LTII refer to irradiation 850 µmol m^-2^ s^-1^ and 1500 µmol m^-2^ s^-1^ respectively. a) c) El Paso 144 (indica cultivar) and b) d) INIA Parao (japonica cultivar). Same letters indicate no significative difference at P < 0.05.

### 3.4 Other important nitrogenous molecules: arginine and GABA accumulation

It is well known that arginine, an important nitrogenous molecule associated with primary nitrogen metabolism in rice, is a metabolite related to urea metabolism, which is highly active in this crop specie. Arginine plays a role in nitrogen storage, nitrogen remobilization and signaling, and as L-amino butyric acid (GABA) and nitric oxide NO precursor (Morris, 2002). For this reason, arginine content was analyzed in response to light and nitrogen content in both cultivars. Results show that El Paso 144 accumulates 3- and 4-times more arginine in leaves with respect to INIA Parao independently of light or nitrogen treatment, indicating a clear difference between the japonica and indica cultivars with respect to arginine metabolism (Fig. 7).

**Fig. 7.**
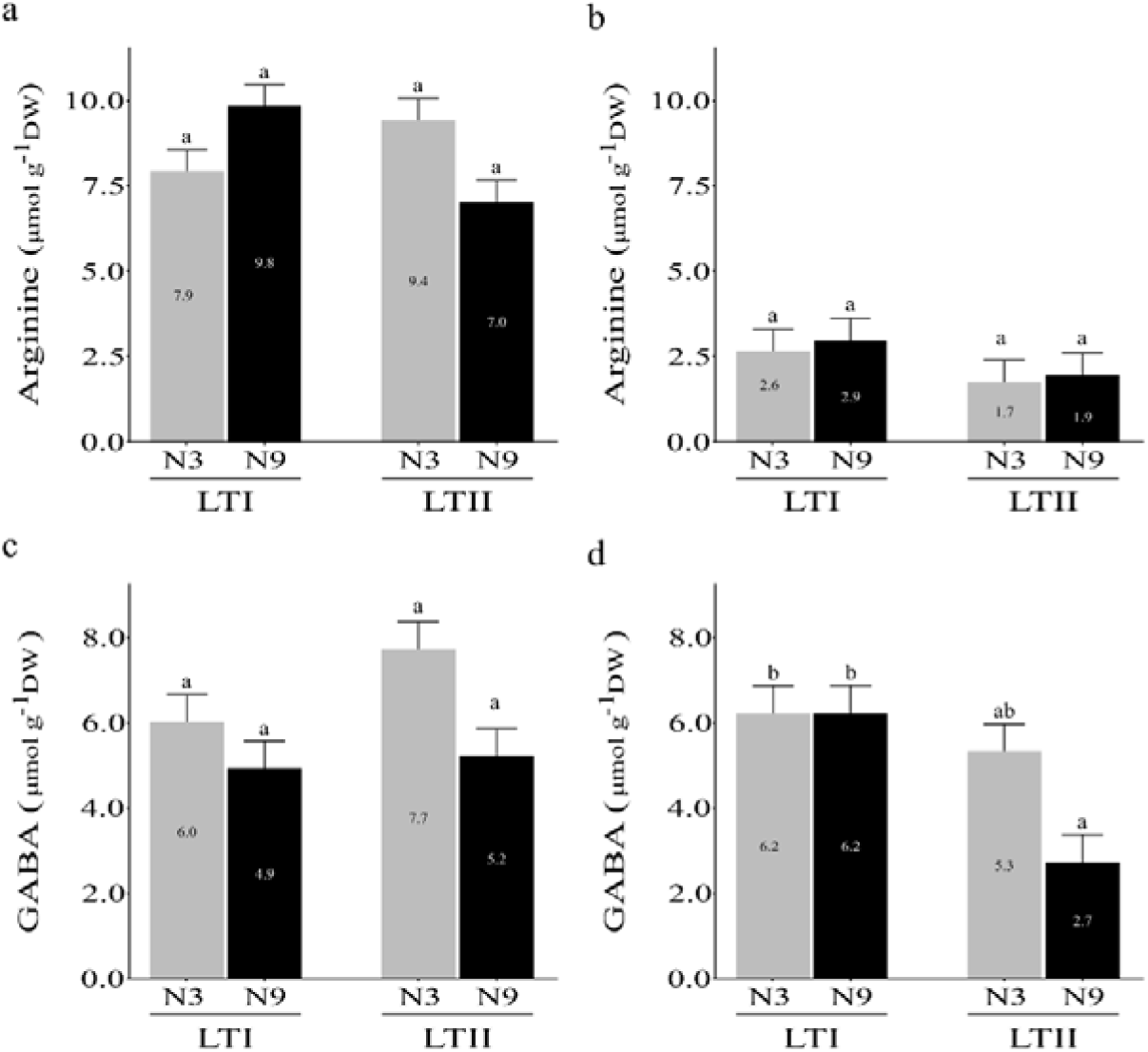
Arginine and L-aminobutyric acid (GABA) in rice leaves. N3 and N9 indicate plants grown under 3 and 9 mM of NO_3_NH_4_, LTI and LTII refer to irradiation 850 µmol m^-2^ s^-1^ and 1500 µmol m^-2^ s^-1^ respectively. a) c) El Paso 144 (indica cultivar) and b) d) INIA Parao (japonica cultivar). Same letters indicate no significative difference at P < 0.05

The content of (GABA), a non-proteic nitrogenous molecule, was analyzed as it plays a role in nitrogen metabolism in response to the environment (Dabravolski and Isayenkov, 2023). The results indicate that the lowest values were found in the high light and high nitrogen treatment. This finding is particularly significant in the japonica cultivar (Fig. 7).

To clear up the complexity of the relationships between GABA, the nitrogenous molecules, and irradiation, a correlation analysis was performed. As depicted in Fig. 8, a negative correlation between GABA and its synthesis intermediates — aspartate, glutamate, and arginine— was observed under low irradiation; this correlation shifts to positive when plants were subjected to high irradiation. This demonstrates a clear re-coordination of the metabolic pathways in response to radiation, resulting in GABA accumulation.

**Fig. 8.**
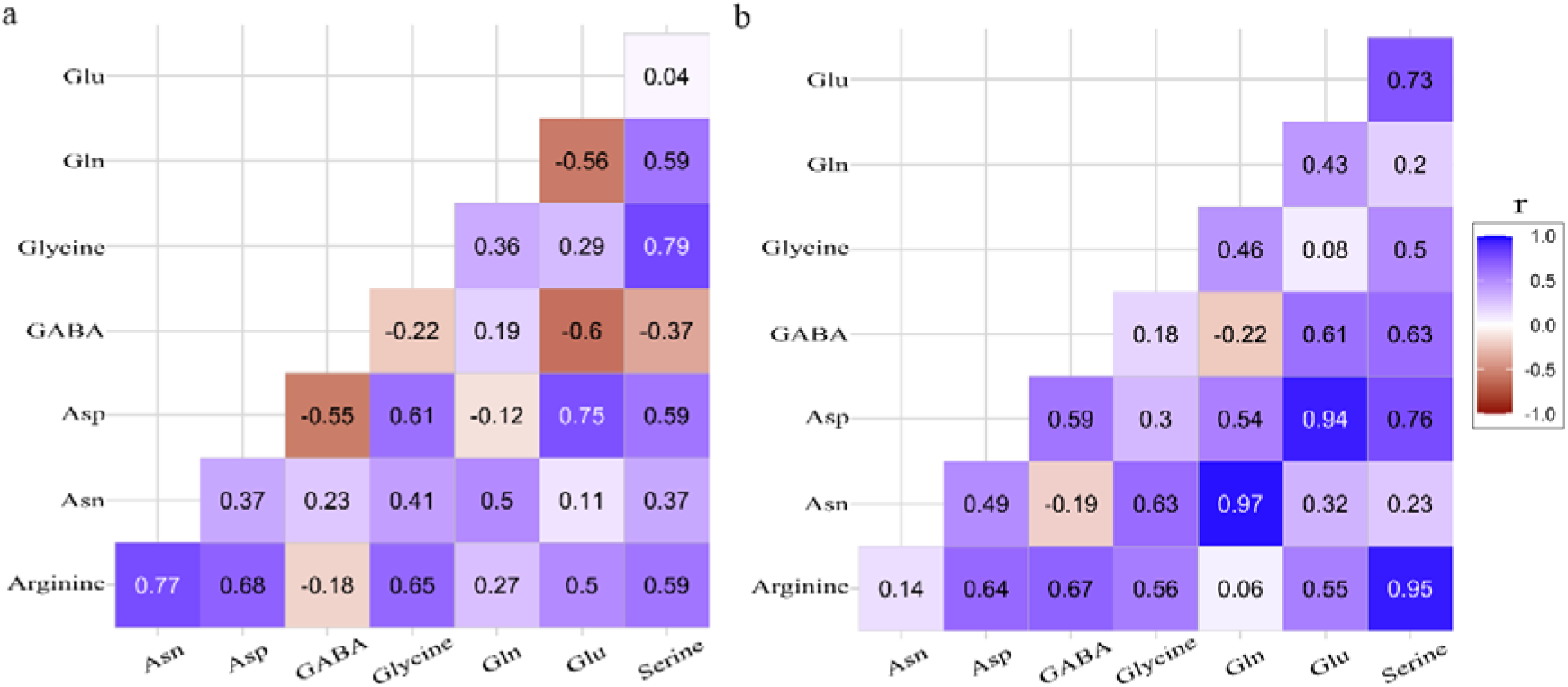
Correlation analysis of the nitrogen metabolites analyzed. a) LTI: Light Treatment I (850 μmol m^-2^ sec^-1^). b) LTII: Light Treatment II (1500 μmol m^-2^ sec^-1^). r: Pearson’s correlation coefficient.

Interestingly, changes in the correlation index in response to the irradiation regimes are also observed in the glutamine and glutamate content, which shifts from -0.56 in LTI to 0.43 as the irradiation increases. This aligns with the previously shown results about changes in the glutamine/glutamate ratio across the different treatments (Fig. 8).

## 4. Discussion

Improving photosynthetic efficiency is widely recognized as one of the most promising strategies to enhance nitrogen use efficiency (NUE) and sustain yield gains in cereals (Thomsen et al., 2014; You et al., 2023). The present study provides physiological evidence supporting this concept by showing that nitrogen metabolism in high-yield temperate rice cultivars is strongly regulated by light irradiance, and that this regulation differs markedly between *indica* and *japonica* subspecies.

Consistent with previous reports, plants supplied with higher nitrogen exhibited increased COL fixation rates, higher leaf N content, and greater accumulation of photosynthetic proteins such as RuBisCO (Makino, 2003). The strong correlation between leaf nitrogen concentration and photosynthetic capacity reflects the predominant allocation of N to chloroplast proteins, particularly those of the Calvin–Benson cycle and the electron transport chain (Salesse-Smith et al., 2025). In both cultivars, high nitrogen availability (N9) resulted in greater tissue N content and enhanced photosynthesis, though the magnitude of this response was modulated by light intensity—highlighting the importance of irradiance in determining NUE. These findings align with field observations, where higher N supply increases leaf N concentration and photosynthetic potential, yet excessive application often leads to diminishing returns and substantial N losses through volatilization or leaching (Ladha et al., 2005).

Interestingly, photochemical parameters (Φ_PSII_, Φ_NPQ_, Φ_NO_) were unaffected by either nitrogen or light, suggesting that in these commercial cultivars the nitrogen and lighting regimes do not affect the mechanisms of capture and canalization of the radiation energy (Table 3).

Another component of the photosynthetic process analyzed was the amount of RuBisCO. Plants grown under high nitrogen supply (N9) exhibited higher levels of the large subunit in both cultivars. A positive correlation was observed between leaf N content and RuBisCO abundance, indicating that increased N allocation to leaves enhances RuBisCO accumulation in rice. This relationship has been previously reported (Makino, 2003; Makino et al., 1984a, 1984b). Consistently, plants under N9 treatment accumulated significantly more N than those grown under low nitrogen supply (N3). These findings reinforce the well-established association between leaf N content, RuBisCO concentration, and COL fixation rate (Salesse-Smith et al., 2025). Moreover, increasing RuBisCO content has been proposed as a strategy to enhance photosynthesis and productivity without compromising nitrogen use efficiency (Salesse-Smith et al., 2025).

A strong association exists between total soluble protein concentration and RuBisCO abundance, reflecting the impact of nitrogen supply on the accumulation of photosynthetic proteins. Field studies in rice have documented a linear correlation between leaf nitrogen content, soluble protein levels, and RuBisCO concentration (Chen et al., 2014). Plants with higher RuBisCO consistently exhibited greater protein concentration per gram of dry weight, underscoring the tight coupling between nitrogen allocation and photosynthetic capacity. Moreover, increases in total protein content under high nitrogen may also reflect temporary N storage in the form of reserve proteins (Mu et al., 2018).

The difference observed in the photosynthetic rate between plants grown in low light radiation and high N dose (LTI_N9) and those grown in high light radiation and high N dose (LTII_N9), seems to be due to differences in the radiation in which the plants were grown and evaluated, since both treatments had a similar amount of RuBisCO. The in vivo activity of RuBisCO is regulated by carbamylation and/or by inhibitor binding (Carmo-Silva et al., 2015). The activation state of RuBisCO responds to light-mediated changes in the redox state of the chloroplast stroma and the phosphorylation potential determined by the ATP/ADP ratio of the stroma via regulation of the ATPase activity of RuBisCO activase (Rca) (Portis et al., 2008). Plants grown under high light radiation are likely to have a higher ATP/ADP ratio, resulting in higher Rca activity; a higher amount of activated RuBisCO would generate a higher CO_2_ fixation rate in plants with similar amounts of RuBisCO.

N remobilization in plants is mainly derived from rapidly degraded proteins in the cytosol of cells through highly regulated intracellular systems such as the proteasome (Nelson et al., 2014; Yanagawa et al., 1999).This process is complemented by groups of proteins located in leaf organelles, such as RuBisCO in chloroplasts, via the autophagy pathway (Nelson et al., 2014; Wada et al., 2015), as well as from membrane structural proteins and stem storage proteins. In rice, up to 80% of grain N comes from protein degradation in mature organs such as leaves and stems (Yoneyama et al., 2016). The results obtained from the content of proteins and RuBisCO show that the increase in soluble proteins in the treatments with high N doses is probably largely due to an accumulation of RuBisCO as reserve protein (Mu et al., 2018).

The analysis of NHLL accumulation in the leaves revealed nondetectable levels of this ion. This could be since a large part of the NHLL is, in rice, efficiently assimilated at the root and leaf level as was reported by several authors (Kusano et al., 2011 and references in). Regarding the accumulation of NOLL, the results show that in the condition of high light radiation and high N dose (LTII_N9) the plants accumulated significantly more NO3^-^compared to the other treatments. The nitrate content in plants is commonly seen because of an imbalance between net absorption and assimilation. It is widely accepted that high light intensity reduces the content of NOLL (Cárdenas-Navarro et al., 1999). The effects of daily light intensity can affect accumulation patterns. The variation of the effects of light intensity can be kept low by selecting appropriate harvest times (Anjana and Iqbal, 2007). In this work, the N nutrition of rice was with NH_4_NO_3_; in rice, the synergistic interaction between these two ions in enhancing growth has been demonstrated. The presence of NOLL in the medium enhances the absorption, accumulation and metabolism of NHLL in the root. In addition, the presence of NHLL decreases the metabolism of NOLL (Kronzucker et al., 1999). The results of the accumulation of NOLL observed in the LTII_N9 may attributed to rice preferentially assimilates NHLL instead of NOLL (Kusano et al., 2011) and demonstrate the capacity of these high yield cultivars to assimilate NHLL in high concentration under the growth light condition evaluated. This preference for NHLL assimilation, common in flooded (reductant) rice systems, has important agronomic implications since maintaining an optimal ion balance in the soil solution can improve NUE and minimize N losses (You et al., 2023).

Amino acids (AA) are central to N metabolism, particularly in the primary assimilation of N in the form of *de novo* synthesized AA, as well as AA derived from protein degradation (Yoneyama et al., 2016). When analyzing the data obtained for total soluble amino acids, it is observed that light influences their content (Fig. 2). Plants grown under low light radiation (LTI) had no significant differences in the amount of AA between N doses. However, plants grown under high light radiation showed significant differences in the amount of AA between N doses for El Paso L144 indica cultivars. Plants grown under LTI had more total soluble AA compared to those grown under LTII. This result shows that the interaction between radiation and N affects the primary and secondary metabolism of nitrogen. In this sense, plants grown under LTI showed symptoms like those grown under a lower dose of N. A study showed that when rice plants grew under different doses of N, the plants with the highest amount of total AA in the plants were those grown under higher doses of N (Chen et al., 2013; Zhang et al., 2017). The AA content in leaves seems to be related to the N demand, since plants with lower amounts of AA have a higher demand for N. Understanding these dynamics could inform site-specific N management strategies, allowing adjustments in fertilizer timing or rate to synchronize N supply with crop demand (Dobermann, 2007). That N demand is probably due to the need for AA for the synthesis of proteins and other nitrogen compounds destined for plant growth. The idea AA is not metabolized when the plants growth at low radiation could be explained by the data reported in a previous study by Quero et al. (2019) wherein it is shown this same level of radiation (850 µmol m^-2^ s^-1^) conditions is not well managed by this high yield elite cultivars analyzed in this study.

The analysis of the transport amino acid ratios shows that only the LTI_N3 growth condition led to a significant increase in the Gln/Glu and Asn/Asp ratios compared to the other treatments. Because it has been proposed that in plants, the Gln generated by the assimilation of NH4^+^ through the GS-GOGAT cycle also functions as a signal molecule regulating metabolism, growth, and development (Yang et al., 2017), the enzymes responsible for their synthesis were analyzed. The analysis of the contents of the enzymes GS and Fd-GOGAT in the leaves of the plants showed differences between the treatments in the quantity of these enzymes that could explain the difference in the Gln/Glu ratio. In the high N dose treatments (N9), the two cultivars presented a greater quantity of Fd-GOGAT, the high content of this enzyme under high nitrogen dose could explain more active transformation of Glu to Gln. However, in LTI conditions, the increase of this enzyme in response to high nitrogen is less significant. The rice OsFd-GOGAT mutant *gogat1* shows only 33% of the total GOGAT enzyme activity in leaves exhibiting chlorosis under natural conditions and a premature leaf senescence (Zeng et al., 2017). Thus, it is not rare to assume that under LTI the induction of Fd-GOGAT by nitrogen is not effective then a process of early senescence in young leaves could be induced (Fortunato et al., 2023).

The quantification results for GS show differences in the amounts of GS isoforms. Treatments with high doses of N (N9) showed a higher amount of chloroplastic GS (GSp) compared to cytosolic GS (GSc), in comparison to plants grown in control doses of N (N3) in both cultivars. However, in the indica cultivar El Paso L144, the increase of GSp in response to nitrogen is less evident. In rice, the chloroplastic GS isoform is abundant in the leaf and is primarily responsible for re-assimilation of ammonium produced from photorespiration in chloroplasts and assimilation in plastids of ammonium deriving from nitrate reduction (Kusano et al., 2011). Furthermore, when serine and glycine, two photorespiratory amino acids, were analyzed, the data would confirm the role of GSp under high light levels because the lowest content of these two amino acids was found in this condition. Results confirm that the idea that 850 umol s^-1^ m^-2^ is insufficient to increase the GSp content and reduce the photorespiratory activity in the high yield cultivars. However, in the indica cultivar, the dependence between GS and photorespiratory amino acids is more evident, showing a clear difference in the response to light and N between both ecotypes.

Moreover, arginine, a nitrogenous compound related to nitrogen use efficiency (Ma et al., 2013), was quantified and a contrasting response between the cultivars was found. Parao, the japonica cultivar, accumulated significantly less arginine than the indica El paso L144 cultivar, independently of the light or nitrogen treatment. Recently, it has been proposed that an active arginine metabolism with a high activity of argininosuccinate lyase determines a less root sensibility to ammonium, increasing the use of this nitrogen source (Xie et al., 2023). Also, this study found that an SNP variant in the gene codifying the enzyme determines that the indica subspecies are more sensitive to ammonium than the japonica subspecies. Thus, the idea of whether this genetic difference between both cultivars could be explaining the difference found in our study is reasonable. In this same line, the correlation analysis among the N compound showed that under low radiation a high concentration of GABA is found simultaneously with a low content of its precursors; interestingly, this relation is inverted under high light conditions. GABA has been reported as a connection molecule between N and carbon metabolism (Dabravolski and Isayenkov, 2023) and could indicate a re-coordination of the metabolism to adjust both metabolism under N and/or C limiting conditions. Taken together, these results provide physiological vases that can be translated into agronomic practices aimed at improving N recovery which help to reduce environmental losses, major challenges in sustainable rice intensification (Erisman et al., 2018; Ladha et al., 2020).

In summary, we found a strong interaction between light and N metabolism in high-yield elite rice cultivars, with a differential response in indica and japonica subspecies that coordinates primary N assimilation with carbon availability. In this sense, the study contributes to the analysis of metabolic strategies involving the accumulation of N compounds under solar irradiation conditions, which is essential for enhancing our understanding of N fertilizer use in elite rice genotypes under field conditions.

## 5. Conclusions

This study demonstrates that N metabolism in rice is highly influenced by light irradiance and that this interaction differs between *indica* and *japonica* subspecies. Higher irradiance enhanced N assimilation, amino acid synthesis, and GOGAT activity, revealing a close coordination between photosynthesis and N utilization. These findings suggest that improving photosynthetic efficiency represents a promising pathway to increase NUE and yield potential in rice, providing a physiological basis for developing management and breeding strategies that optimize light interception and N use under field conditions.

## CRediT authorship contribution statement

Gastón Quero: Conceptualization, Writing – review & editing, Validation, Methodology, Formal analysis, Data curation. Pedro Silva: Writing – original draft, Validation, Methodology, Investigation. Maria Martha Sainz: Writing – review & editing, Methodology. Sebastián Fernández: Resources, Methodology. Sebastián Simondi: Software, Formal analysis, Data curation. Jesus Castillo: Funding acquisition, Writing – review & editing. Omar Borsani: Conceptualization,Writing – review & editing, Supervision, Funding acquisition.

## Acknowledgements

The authors gratefully acknowledge Dr. Pedro Díaz Gadea for his valuable contributions to the promote the idea of this work and provide his scientifically background to enrich the discussion.

