## Supplemental data for "Irradiation and nitrogen metabolism: differential responses in high yield indica and japonica rice commercial cultivars"

**Journal of Plant Physiology**

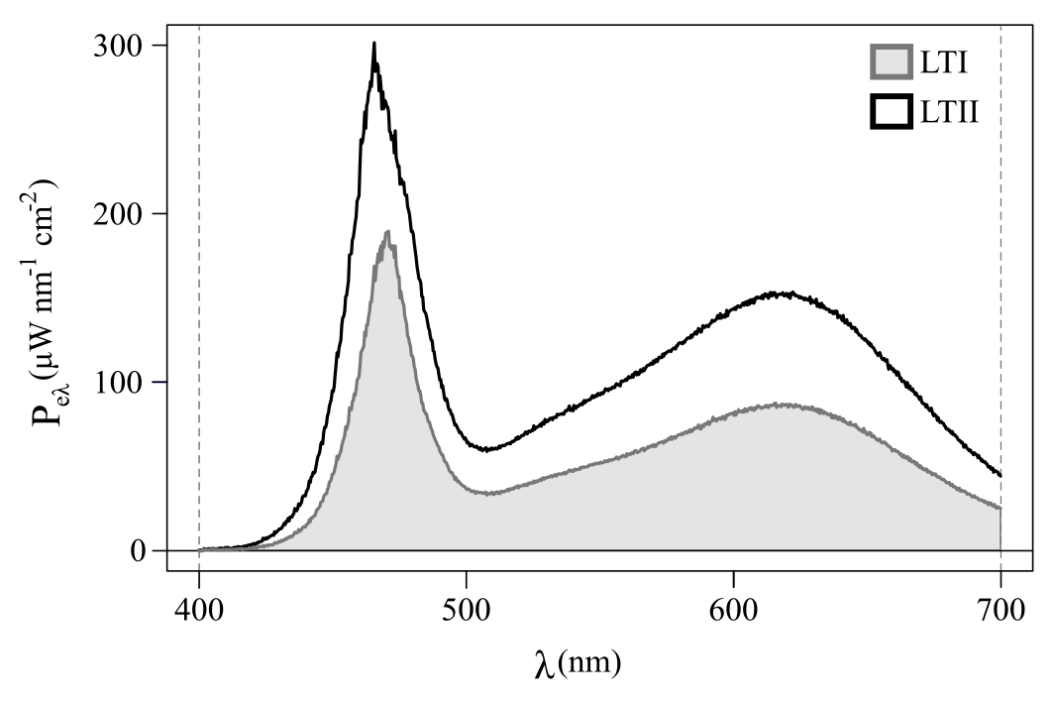

**Fig. 1S.** Spectral distributions of the two light treatments depicted as Spectral Power (P_eλ_) as a function of wavelength (λ). LTI: Light Treatment I (850 μmol m^-2^ sec^-1^). LTII: Light Treatment II (1500 μmol m^-2^ sec^-1^).

Table SI Nutrient solution Yoshida et al. 1976

| Macronutrients | mM |
| --- | --- |
| MgSO_4_:7H_2_O | 1.6 |
| CaCl_2_ | 1.0 |
| NaH_2_PO_4_.2H_2_O | 0.32 |
| K_2_SO_4_ | 0.5 |
| Micronutrients | μM |
| Citric acid monohydrate | 70.8 |
| FeCl_3_·6H_2_O | 35.4 |
| MnCl_2_·4H_2_O | 9.5 |
| H_3_BO_3_ | 19.0 |
| CuSO_4_.5H_2_O | 0.15 |
| ZnSO_4_.7H_2_O | 0.15 |
| (NH_4_)_6_·MO_7_O_24_·4H_2_O | 0.07 |

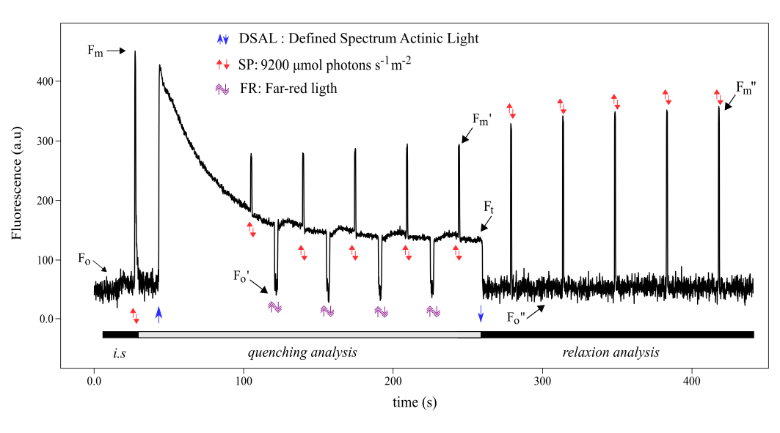

**Fig. 2S** Fluorescence quenching analysis followed by relaxation analysis using modulated fluorescence. Blue up and down arrows indicate that light is turned on and off, respectively. Red arrows indicate the position of saturating pulses (SP). Double head arrow indicates a far-red light pulse (FR).
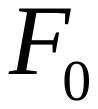
,
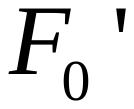
, and
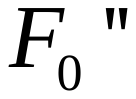
represent minimum fluorescence in the initial phase, quenching analysis phase, and relaxation analysis phase, respectively.
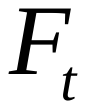
 is the currente fluorescence for light-adapted states in the time t;
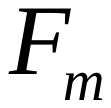
and
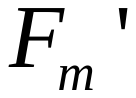
 represent maximum fluorescence in dark and light conditions, respectively.
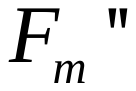
 is the maximum fluorescence in dark conditions during recovery.

**Table 2S** Glutamine synthetase enzyme activities.

| Genotype | Light | Nitrogen | GS  (U mg^-1^) |
| --- | --- | --- | --- |
| El Paso 144 | LTI | N3 | 106 a |
|  |  | N9 | 81 a |
|  | LTII | N3 | 145 a |
|  |  | N9 | 101 a |
| INIA Parao | LTI | N3 | 117 a |
|  |  | N9 | 127 a |
|  | LTII | N3 | 151 a |
|  |  | N9 | 93 a |

U mg^-1^: activity units per mg of protein.
